# Age-related dysregulation of the retinal transcriptome in African turquoise killifish

**DOI:** 10.1101/2024.02.21.581372

**Authors:** Steven Bergmans, Nicole C. L. Noel, Luca Masin, Ellen G. Harding, Aleksandra M. Krzywańska, Julie D. De Schutter, Rajagopal Ayana, Chi-Kuo Hu, Lut Arckens, Philip A. Ruzycki, Ryan B. MacDonald, Brian S. Clark, Lieve Moons

## Abstract

Age-related vision loss caused by retinal neurodegenerative pathologies is becoming more prevalent in our ageing society. To understand the physiological and molecular impact of ageing on retinal homeostasis, we used the short-lived African turquoise killifish, a model known to naturally develop central nervous system (CNS) ageing hallmarks and vision loss. Bulk and single-cell RNA-sequencing (scRNA-seq) of three age groups (6-, 12-, and 18-week-old) identified transcriptional ageing fingerprints in the killifish retina, unveiling pathways also identified in the aged brain, including oxidative stress, gliosis, and inflammageing. These findings were comparable to observations in ageing mouse retina. Additionally, transcriptional changes in genes related to retinal diseases, such as glaucoma and age-related macular degeneration, were observed. The cellular heterogeneity in the killifish retina was characterised, confirming the presence of all typical vertebrate retinal cell types. Data integration from age-matched samples between the bulk and scRNA-seq experiments revealed a loss of cellular specificity in gene expression upon ageing, suggesting potential disruption in transcriptional homeostasis. Differential expression analysis within the identified cell types highlighted the role of glial/immune cells as important stress regulators during ageing. Our work emphasises the value of the fast-ageing killifish in elucidating molecular signatures in age-associated retinal disease and vision decline. This study contributes to the understanding of how age-related changes in molecular pathways may impact CNS health, providing insights that may inform future therapeutic strategies for age-related pathologies.

**Highlights:**

- The aged killifish retina displays several ageing hallmarks, such as oxidative stress, gliosis and inflammageing, at the transcriptome level.
- Risk genes for neurodegenerative disorders show dysregulation in the old killifish retina.
- All vertebrate retinal cell types are present in the killifish retina.
- Transcriptional dysregulation in the aged killifish retina is observed across cell types.

## 1| Introduction

The mammalian central nervous system (CNS) is vulnerable to the accumulation of age-related pathologies that lead to progressive, irreversible disease. Ageing is also a major risk factor for many retinal neurodegenerative diseases that severely impact visual function and quality of life (Congdon, 2004; Enoch et al., 2019; Swenor et al., 2020). Neurodegenerative diseases affecting the light-sensing retina, including glaucoma and age-related macular degeneration (AMD), are becoming increasingly prevalent as the life expectancy increases (Klein and Klein, 2013). There are currently no treatments available to restore vision after retinal neurodegeneration, nor targeted therapies to prevent degeneration and related vision loss in the long-term. Therefore, novel insights into retinal ageing are valuable to determine the physiological and molecular consequences of ageing on retinal homeostasis and identify mechanisms underpinning age-related retinal dysfunction.

Small animals such as mice, rats, and zebrafish have been extensively utilised to study fundamental mechanisms underlying different retinal neurodegenerative diseases (Collin et al., 2020; Koh et al., 2019; Niwa et al., 2016; Noel et al., 2022; Veys et al., 2021). Unfortunately, ageing studies are challenging in mammalian model organisms due to their relatively long lifespan (Kim et al., 2016). Zebrafish naturally undergo age-related retinal changes that include photoreceptor degeneration, gliosis, and vision loss (Martins et al., 2022; Van houcke et al., 2019). However, like mice and rats, zebrafish also have relatively long lifespans, and can live up to 5 years (Kim et al., 2016). Invertebrate gerontology models such as *Caenorhabditis elegans* or *Drosophila melanogaster,* which have short lifespans of up to a few weeks, do not allow for comparable study of the aged retina as *C. elegans* lack a visual system and the *Drosophila* retina differs from the vertebrate retina in terms of structure and cell types. This demonstrates the pressing need for a genetically tractable vertebrate model organism with conserved retinal organisation and a condensed lifespan, facilitating high-throughput research into mechanisms underlying retinal ageing.

The short-lived African turquoise killifish (*Nothobranchius furzeri*) is emerging as a leading vertebrate genetic model system for studies of ageing (Kim et al., 2016; Moses et al., 2023; Platzer and Englert, 2016). The inbred GRZ strain has the shortest lifespan in captivity amongst vertebrates, with a median life expectancy between four and six months, and rapidly presents ageing phenotypes (Kim et al., 2016; Van houcke et al., 2021; Vanhunsel et al., 2021). Several short-lived killifish strains have been described to show macroscopic ageing traits similar to those observed in humans, such as depigmentation, spinal curvature, muscle atrophy, reduced locomotor function, and cognitive decline upon ageing (Cellerino et al., 2016; Kim et al., 2016; López-Otín et al., 2013; Mariën et al., 2024; Valenzano et al., 2015), as well as molecular hallmarks of human ageing, including oxidative stress, cellular senescence, altered cellular communication, dysregulated nutrient sensing, and stem cell exhaustion (Baumgart et al., 2014; Valenzano et al., 2006).

Our recent characterisations of the killifish visual system identified the presence of several ageing hallmarks in the retina and optic tectum — the main area of the brain that performs visual processing in teleost fish — including increased levels of oxidative stress, DNA damage, altered cellular communication, inflammageing, stem cell depletion, and increased expression of cellular senescence markers (Vanhunsel et al., 2021). Interestingly, like the killifish brain (Bagnoli et al., 2022; Matsui et al., 2019), the killifish retina presented signs of thinning and neurodegeneration upon ageing. These ageing characteristics were associated with a marked decline in visual acuity (Vanhunsel et al., 2021).

We performed unbiased transcriptomics profiling to better understand the molecular signatures of retinal ageing in the killifish. We first assayed the retinal transcriptome dynamics across the ageing process using bulk RNA-sequencing (RNA-seq), identifying transcriptional shifts from young adult (6-week-old) to old (18-week-old). We investigated the cell type specificity of ageing phenotypes using single-cell RNA-sequencing (scRNA-seq), defining retinal cell populations and identifying age-induced changes in transcript levels within individual retinal cell types. We determined that there is ageing-associated transcriptome dysregulation across all retinal cell populations, highlighting systemic changes in transcriptional homeostasis that likely contribute to reduced killifish retinal function with age. From these studies, we highlight the utility of killifish for understanding the molecular aetiology of retinal ageing.

## 2| Results

### 2.1| Bulk RNA-seq analysis identifies age-related transcriptional shifts in the killifish retina

We first profiled retinal transcript expression using bulk RNA-seq to characterise transcriptional alterations that may underly the killifish ageing phenotypes. We investigated young adult (6-week-old), middle aged (12-week-old), and old (18-week-old) female fish, using 10 biological replicates for each age group (Figure 1A). Whole neural retinas were dissected from the eye after retinal pigmented epithelium (RPE) removal and subsequently processed for bulk RNA-seq. Principal component analysis revealed a gradual shift in transcriptional state that correlated with age (Figure 1B). We identified 65 and 401 up-regulated genes between 6- and 12-week-old, and between 6- and 18-week-old retinas, respectively, by direct comparison of gene expression across individual ages (cut-offs: FDR < 0.05, |log2FC| ≥ 1). Of these differentially expressed transcripts, 59 were shared across both comparisons (Figure 1C). A smaller number of transcripts showed decreased expression across retinal ageing, with 17 genes having decreased transcript expression between 6- and 12-week-old and 83 genes between 6- and 18-week-old retinas, of which 12 genes were shared across age comparisons (log2FC < 1 & FDR < 0.05; Figure 1C). A heatmap and table including all differentially expressed transcripts across killifish retinal ageing are provided in Figure S1 and Table S1. We focused on genes changing in expression between 6- and 18-week-old samples for subsequent analyses, as these data highlighted the greatest age-associated transcriptional disparity.

**Figure 1.**
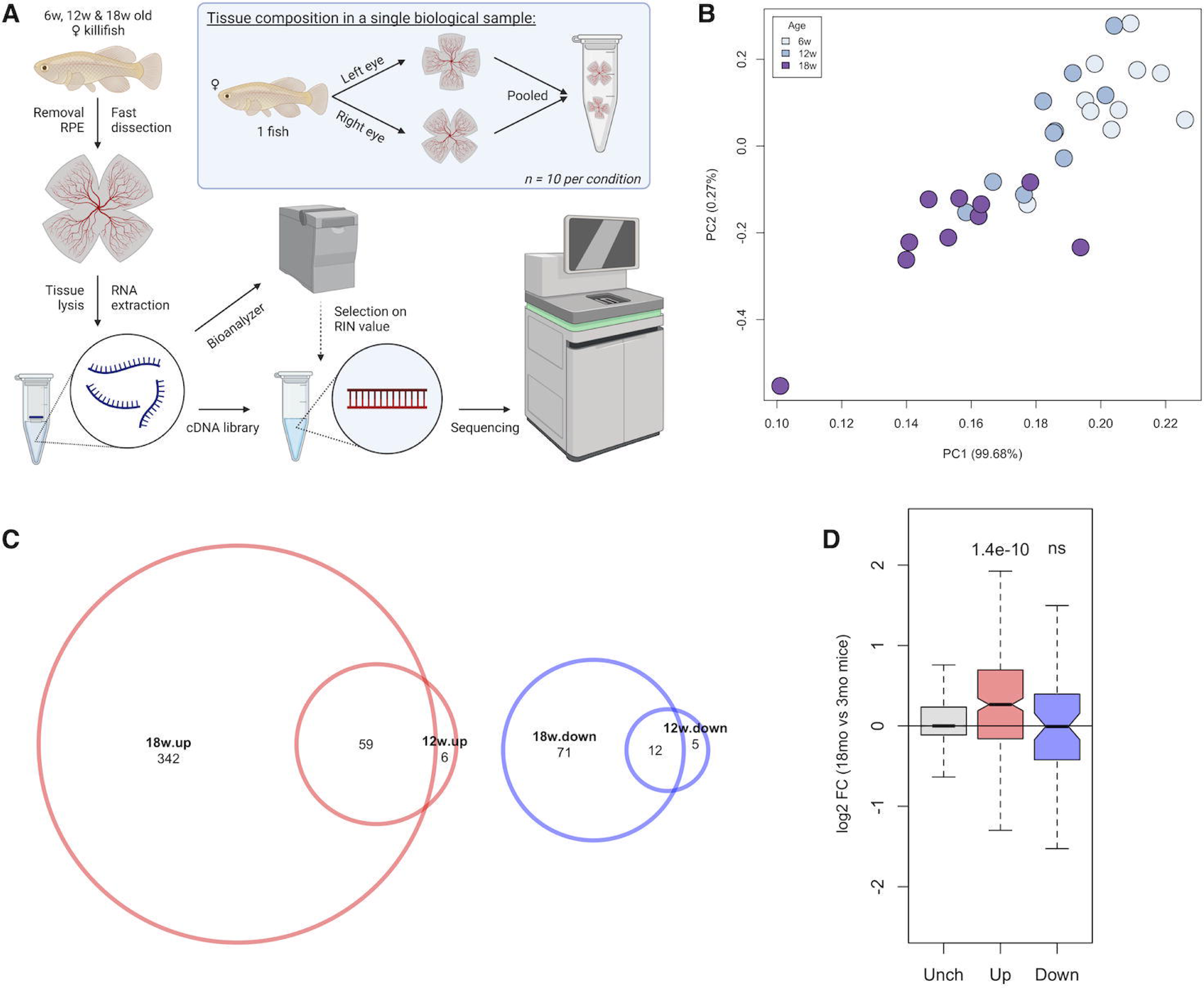
Turquoise killifish retinas have age-related changes in gene expression. (A) Experimental setup for the bulk RNA-seq experiments. (B) Principal component analysis of 6-, 12-, and 18-week-old individual RNA-seq samples (n=10 each) showing that age-related variance in transcriptome across samples. (C) Scaled Venn diagrams depicting up- (red) and down-regulated (blue) genes between young (6-week-old), middle-aged (12-week-old) and aged (18-week-old) retinas. There are more up-regulated genes than down-regulated genes identified in the dataset with age. Differentially expressed genes were identified using following thresholding criteria: FDR < 0.05 and |log2FC| ≥ 1. (D) Boxplots representing the age-dependent fold change of transcript expression of mouse orthologs (Xu et al., 2022) to Up, Down, and Unchanged sets of differentially expressed killifish genes. The killifish-specific unchanged (grey) and up-regulated genes (red) show a similar trend of expression as in the ageing mouse retina. This is not observed for the down-regulated genes (blue) (Wilcoxon rank-sum test). FC = fold change, mo = months, ns = not significant, PC = principal component, RIN = RNA integrity number, RPE = retinal pigment epithelium, Unch = unchanged, w = weeks.

We next compared our dataset to published mouse ageing retinal transcriptomes (3-month-old versus 18-month-old (Xu et al., 2022)) to investigate the transcriptional similarities between killifish and mammalian retinal ageing. Genes with increased expression during killifish retinal ageing showed a similar trend to their mouse orthologs (Figure 1D). In contrast, genes that decreased in the ageing killifish dataset did not show consistency with the mouse findings. As the genes we identified as down-regulated in killifish retina included many retinal pigment epithelium (RPE)-related genes, such as *rpe65a*, *tyr*, and *pmelb* (Table S1), this may indicate some down-regulated genes are the technical artifact of differences in RPE removal between young and old killifish. Younger killifish have higher adherence of the RPE to the neural retina than old killifish, and RPE contamination may therefore be higher in the dataset for the young age group. Gene ontology (GO) term analysis of our up-regulated gene dataset revealed pathway enrichments related to oxidative stress (GO:0051775, GO:0034599, GO:0006979), metabolism (GO:0042572, GO:0031325), and inflammation (GO:0090025, GO:0043368); these findings coincide with those of the aged murine retinal transcriptome for pathways correlated to oxidative stress and inflammatory responses (Xu et al., 2022), which had age-related changes in similar categories. A detailed list containing the GO categories of all up- and down-regulated biological processes (BP) and molecular functions (MF) of the aged killifish is included in Table S2.

We sought to first validate important processes associated with CNS ageing as well as killifish retinal ageing hallmarks that were previously identified using histological analyses (Mattson and Arumugam, 2018; Vanhunsel et al., 2021). For this, we looked at transcripts for genes involved in oxidative stress, gliosis, inflammageing, and cellular senescence. Oxidative stress occurs due to the accumulation of reactive oxygen species and is a prominent driver of progressive loss of tissue function during ageing (Liguori et al., 2018). We observed increased transcript levels for the oxidative stress response genes *pdk4* (log2FC = 1.16, p-value = 4.98E-12, FDR = 1.81E-10) and *slc7a11* (log2FC = 1.48, p-value = 6.25E-15, FDR = 3.62E-13) in the aged killifish retina (Figure 2A) (Gao et al., 2022; Yan et al., 2023). Furthermore, we detected increased transcript levels for the genes *tgfb3* (log2FC = 1.04, p-value = 1.54E-11, FDR = 5.10E-10) and *rlbp1a* (log2FC = 1.03, p-value = 3.31E-15, FDR = 2.05E-13), known to be related to gliosis (*tgfb3*) (Conedera et al., 2021) and Müller glia homeostasis (*rlbp1a*) (Vázquez-Chona et al., 2009) (Figure 2A). Glia become reactive in response to stress or injury, a process referred to as gliosis that is defined by morphological and gene expression changes (Bringmann et al., 2009; Seitz et al., 2013). Using immunohistochemical labelling for Rlbp1, we correlated the increased *rlbp1a* transcript expression with an appreciable rise in immunofluorescent signal in aged retinas (Figure 2B). Rlbp1 immunostaining also revealed signs of expansion and elaboration of Müller glia morphology in old fish (Figure 2B), which was confirmed by immunolabeling for the canonical Müller glia marker glutamine synthetase (GS) (Figure 2C). Further, the aged immune system is known to undergo inflammageing, where it alters cell number, morphology, and secretory profile (Godbout and Johnson, 2009; Vanhunsel et al., 2021). Our bulk RNA-seq data revealed an up-regulation of inflammageing markers such as *nfkb2* (log2FC = 1.07, p-value = 4.78E-10, FDR = 1.174E-8) and *apoeb* (log2FC = 1.99, p-value = 8.01E-49, FDR = 6.09E-46) in the old retina (Figure 2A) (García-García et al., 2021; Godbout and Johnson, 2009). Morphological assessments using *in situ* hybridization chain reaction (HCR) for *apoeb* highlighted a visible increase in the amount of *apoeb* in the aged retina as well as differences in morphology between 6 and 18 weeks (Figure 2D). We confirmed that *apoeb* labelled immune cells with antibody co-labelling for the pan-leukocyte marker L-plastin (Figure 2E). Thus, we observed increased oxidative stress levels as well as Müller glia and microglia reactivity in the aged killifish retina, consistent with previous reports (Vanhunsel et al., 2021). However, unlike our previous study characterizing senescence markers (*cdkn1a/*p21 and *cdkn1b/*p27) by quantitative reverse-transcription PCR, we did not observe significant transcriptional changes for senescence-related genes between young and old neural retinas in the bulk RNA-seq dataset (Figure 2A).

**Figure 2.**
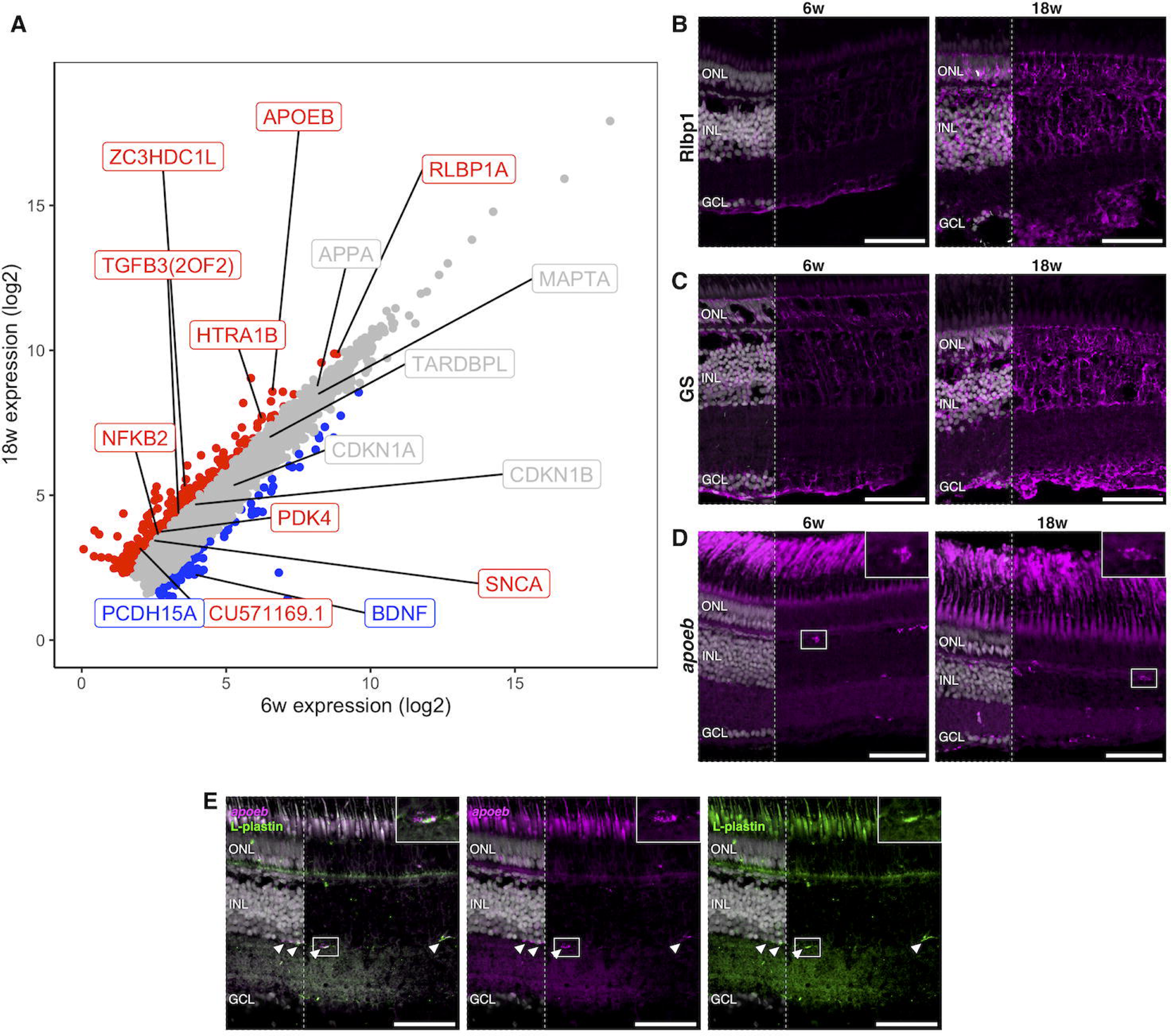
Aged killifish retinas show signs of gliosis, inflammageing, and neurodegeneration detected by bulk RNA-seq. (A) Scatter plot showing mean expression of all genes detected by bulk RNA-seq across 6w and 18w samples. The different colours denote genes that passed FDR and FC thresholds (FDR < 0,05 and|log2FC| ≥ 1). Genes that are not significantly changed are shown in grey; up-regulated genes are in red; down-regulated genes in blue. ZC3HDC1L is representing the *optn* gene and CU571169.1 is *slc7a11.* (B) Immunohistochemistry for Rlbp1 shows a visual increase in Rlbp1 protein expression in aged retinas, corresponding to increased transcript expression in RNA-seq. Additionally, images highlight Müller glia expansion and elaboration. (C) Immunostaining of glutamine synthetase similarly shows expansion of Müller glia cell morphology in 18-week-old killifish retinas, as compared to 6-week-old retinas. (D) *in situ* HCR for *apoeb* in 6 and 18-week-old killifish retinas highlight an increase in immune cells (microglia/macrophages) with age as well as changes in their morphology (insets), which are signs of inflammageing. (E) Microglia/macrophage-specific labelling by *apoeb* is confirmed using antibody co-labelling with the pan-leukocyte marker L-plastin. Arrowheads indicate cells that are both *apoeb+* and L-plastin+. Images are acquired as mosaic Z-stack and are visualised as maximum projections. Merged images with nuclei shown in left third of image; remaining image is without nuclei. Scale bars = 50 µm. HCR = hybridisation chain reaction, FC = fold change, FDR = false discovery rate, GCL = ganglion cell layer, GS = glutamine synthetase, INL = inner nuclear layer, ONL = outer nuclear layer, unch = unchanged, w = weeks.

We next investigated whether retinal degeneration-associated genes were changing across ageing in the killifish transcriptome. While pathogenic *OPTN* variants are known to result in hereditary forms of glaucoma (Rezaie et al., 2002), increased *OPTN* levels are associated with neuroprotective processes (Markovinovic et al., 2018; Weil et al., 2018). Our dataset revealed an age-related increase in *optn* transcript levels (log2FC = 1.79, p-value = 4.36E-33, FDR = 1.43E-30; Figure 2A), but the effect of this increase remains elusive. Brain-derived neurotrophic factor (BDNF) is important for neuronal homeostasis and survival (Chiavacci et al., 2022; Lima Giacobbo et al., 2018). We observed a decrease in *bdnf* (log2FC = −1.87, p-value = 1.20E-26, FDR = 2.66E-24) expression with age (Figure 2A), which may result in a heightened risk of developing retinal pathology. Indeed, BDNF is of interest for treating neurodegenerative diseases such as glaucoma and is reduced in glaucomatous eyes (Gupta et al., 2014; Osborne et al., 2018). Variants in the *HTRA1* promoter and gene increase the risk for developing AMD (DeAngelis et al., 2008; Friedrich et al., 2015; Tam et al., 2008; Yang et al., 2006). Interestingly, Htra1 protein levels increased with age in the zebrafish retina and elevated *htra1* transcript levels were observed in a zebrafish photoreceptor degeneration model (Oura et al., 2018). The expression of *htra1b* increased (log2FC = 1.40, p-value = 1.86E-25, FDR = 3.71E-23) in the aged killifish retina (Figure 2A), which may be indicative of photoreceptor dysfunction. Pathogenic variants in *PCDH15* cause Usher syndrome, a condition characterised by hearing and vision loss (Ahmed et al., 2001). *pcdh15a* expression decreased (log2FC = −1.11, p-value = 7.06E-8, FDR = 9.25E-7) with age in the killifish retina (Figure 2A), which may also suggest degradation of photoreceptor health. Thus, genes known to be associated with retinal degeneration show age-dependent expression changes in the killifish retina, highlighting the value of the killifish retina as an ageing model.

We next examined changes in transcript levels of genes associated with degenerative disease, as pathological hallmarks of neurodegenerative disorders can also manifest within the retina (London et al., 2013; Veys et al., 2019). Interestingly, we observed increased transcript levels of the gene encoding alpha-synuclein (*snca*), known to be a key player in Parkinson’s disease and up-regulated in the killifish brain (Matsui et al., 2019), in the aged killifish retina (log2FC = 1.13, p-value = 1.28E-10, FDR = 3.51E-9; Figure 2A). In contrast, we did not observe differential expression of genes related to other neurodegenerative diseases, such as Alzheimer-related genes *mapta* (encoding tau protein) or *appa* (encoding amyloid beta precursor protein) and the frontotemporal dementia/amyotrophic lateral sclerosis gene *tardbp* (encodes Tdp-43 protein). This could indicate that features reported in the aged killifish brain, such as amyloid beta-plaques and Tdp-43 stress granules (Bagnoli et al., 2022; Bergmans et al., 2023a; de Bakker and Valenzano, 2023; Louka et al., 2022; Matsui et al., 2019), are not manifesting in the ageing killifish retina, although investigation at the protein level would be required to determine this.

In conclusion, bulk RNA-seq analysis of the aged killifish retina identified conserved age-related changes in genetic pathways associated with retinal ageing and neurodegenerative disease.

### 2.2| scRNA-seq catalogues all retinal cell types in the killifish retina

Age-associated retinal diseases, including AMD and glaucoma, are known to selectively impact individual cell types leading to their progressive loss, i.e., photoreceptors and retinal ganglion cells, respectively. This is likely the result of a complex interplay between genetics and the environment, as disease risk-associated variants are not wholly restricted to genes expressed only in the cell type(s) primarily affected by disease pathology and can underlie different diseases in a variant-specific manner. For instance, *OPTN* is not only expressed in retinal ganglion cells, but is also found in a subset of retinal interneurons, the brain, and non-neural tissues such as the heart (Kroeber et al., 2006; Rezaie and Sarfarazi, 2005). However, glaucoma-associated disease variants lead specifically to retinal ganglion cell death (Rezaie et al., 2002). One hypothesis is that the ageing process and related stresses causes asymmetric alterations in gene regulation across cell types. Alternatively, ageing may manifest as a more general process, resulting in global changes in gene expression across all cell types, with environmental factors contributing to disease onset and/or severity. To investigate how gene expression changes across retinal ageing affects transcript expression within individual cell types, we performed scRNA-seq of the killifish retina age-matched to the bulk RNA-seq experiments (Figure 3A). 10x Genomics 3’ sequencing was performed on two biological replicates at each age (6-, 12-, and 18-week-old animals), capturing 37,243 cells across the ages with an average transcript count of 3,311 ± 2,672 per cell (Figure S2).

**Figure 3.**
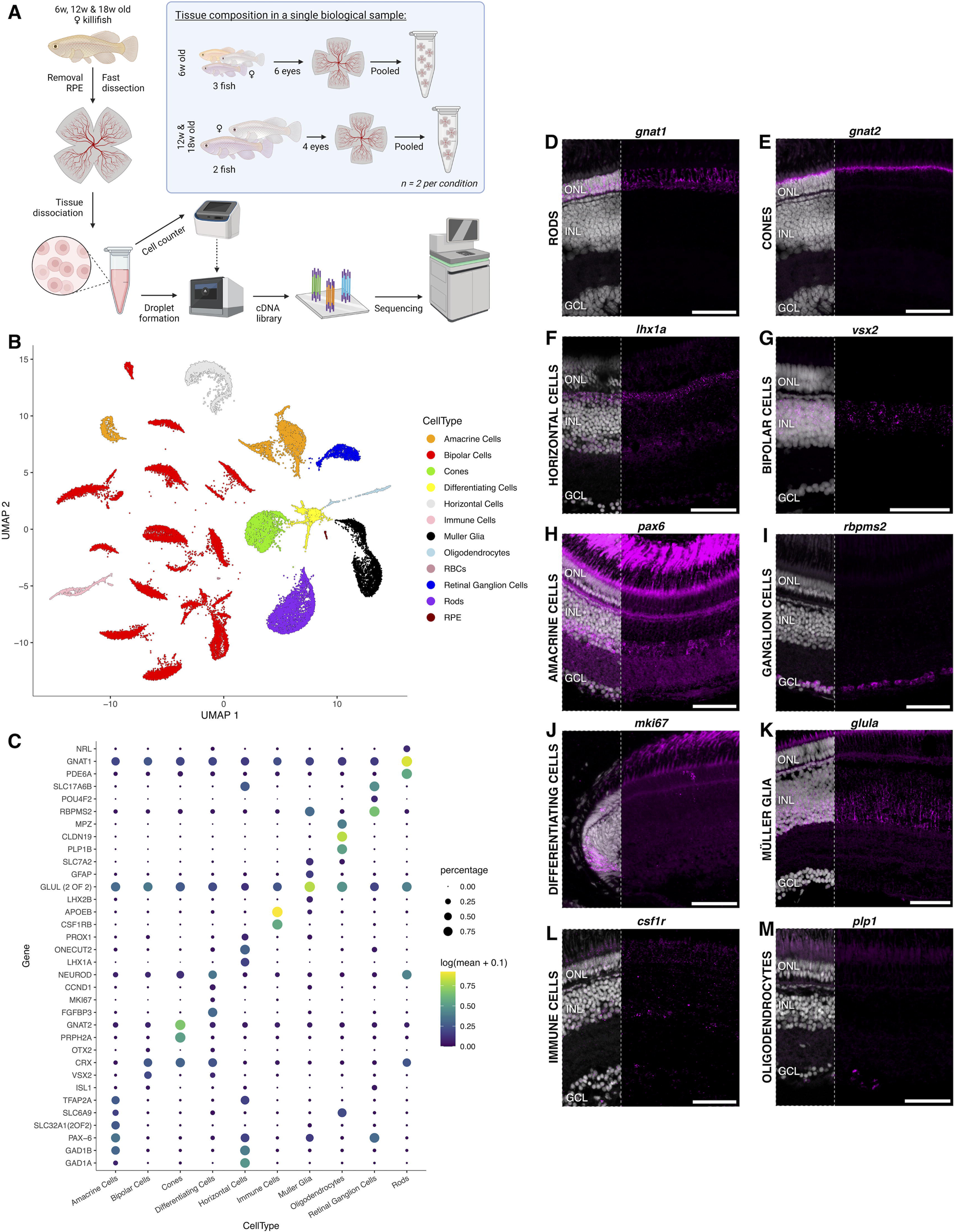
scRNA-seq identifies neuronal and glial cell types within the killifish retina. (A) Experimental setup for the single-cell RNA-sequencing experiments. (B) UMAP dimension reduction of the killifish retina scRNA-seq dataset with clusters coloured by annotated retinal cell type. All cell types expected of a vertebrate retina, as well as oligodendrocytes, are present in the killifish retina. (C) Dot plot showing the specificity of marker genes within individual retinal cell types. The size of the dot represents the percentage of cells within the population expressing transcripts for the gene, while the colour indicates the average expression across individual cells. (D-M) Spatial validation of cell type marker genes via *in situ* HCR confirms the cell type identification for every retinal cell type. Images are acquired as mosaic Z-stack and are visualised as maximum projections. Merged images with nuclei shown in left third of image. Scale bars = 50 µm. HCR = hybridisation chain reaction. UMAP = uniform manifold approximation and projection. GCL = ganglion cell layer, INL = inner nuclear layer, ONL = outer nuclear layer, RBC = red blood cell. RPE = retinal pigment epithelium, w = weeks.

Dimension reduction resulted in discrete clustering of cell populations (Figure 3B, Figure S2B). Each cluster was comprised of cells from each biological replicate and timepoint except for clusters 23 and 29 (28 and 27 cells, respectively; Figure S2C). Known marker genes for major cell classes were used to distinguish retinal cell types, including cone photoreceptors (*gnat2*), rod photoreceptors (*nrl*, *rho*), horizontal cells (*lhx1a*), bipolar cells (*vsx2*), Müller glia (*lhx2b*, *aqp1a.1*), amacrine cells (*tfap2b*, *pax6*), and retinal ganglion cells (*pou4f2*, *gap43*) (Figure 3B-C, Figure S3). We also observed clusters containing immune cells (microglia/macrophages; *aif1*, *c1qc*), red blood cells (Nfu_p_1_007730, encoding haemoglobin subunit alpha A), and RPE cells (*rpe65*) (Figure S4), as well as a cluster containing cells marked by proliferative markers (*ccnd1*) and markers of retinal neurogenesis (*neurod*; differentiating cells) (Figure 3B, Figure S3). All cell type identities were maintained across ageing, and while there may be shifts in cell proportion between the samples investigated, we did not observe substantial age-dependent loss of individual cell types (Figure S2D). Previous morphological studies on killifish retina found that there was an age-related increase in immune cells and a decrease in stem cells/differentiating cells (Vanhunsel et al., 2021). The cell proportions estimated from the scRNA-seq results coincide with these findings (Figure S2E) – however, capture efficiency biases of individual cell types cannot be ruled out.

Cell type cluster markers (neurons, glia, immune cells) were validated on tissue using *in situ* HCR (Figure 3D-M, Figure S5A-I), confirming the basic vertebrate retinal architecture. Briefly, the retina is comprised of three nuclear layers: the outer nuclear layer (ONL), inner nuclear layer (INL), and ganglion cell layer (GCL), separated by plexiform (synaptic) layers, namely outer and inner plexiform layers (OPL and INL, respectively) (Yamagata et al., 2021). General photoreceptor markers (*crx*) (Figure S5A) and photoreceptor subtype-specific labelling for rods (*gnat1, nrl)* (Figure 3D, Figure S5B) and cones (*gnat2*) (Figure 3E) were confined to the outer retina, with rod nuclei sitting below cone nuclei. Horizontal (*lhx1a, tfap2a*) (Figure 3F, Figure S5C), bipolar (*vsx2, otx2*) (Figure 3G, Figure S5D), and amacrine (*pax6, slc32a1b*) (Figure 3H, Figure S5E) cells were observed in the INL and localized to their specific sublayers: the apical INL, dispersed throughout the INL, and along the basal INL, respectively. Retinal ganglion cells in the GCL were labelled with markers *rbpms2* and *slc17a6b* (Figure 3I, Figure S5F). Differentiating cells, marked by *stmn1a, fgfb3*, and *mki67* (Figure 3J, S5G), were present in the ciliary marginal zone, the neurogenic niche at the retinal periphery. Müller glia (*glula, rlbp1a*) showed characteristic radial fibre morphology with their nuclei confined to the INL (Figure 3K, Figure S5H), while *gfap* signal was restricted to their end feet (Figure S5H). The resident immune cells, marked by *csf1r*, lined the IPL (Figure 3L), as expected for resting microglia/macrophages (Murenu et al., 2022). Notably, in contrast to mammals, we identified oligodendrocytes (*plp1b, cldn19*), highly abundant in the optic nerve head (Figure S5I) and sparsely distributed throughout the retina in the GCL (Figure 3M, Figure S5I) and the INL (Figure S5I, arrowhead) (Nakazawa et al., 1993; Santos-Ledo et al., 2023).

For retinal cell types showing heterogeneous clustering in the UMAP, we also identified and validated markers that distinguished subtypes when subclustering resolution permitted. We further investigated the expression of subcluster marker genes in the other identified cell types (Figure S6). We subclustered cone photoreceptors into L/M, S, and UV cones (Figure 4A, B). L and M cones are often physically fused as double cones in fish species (Siebeck et al., 2014), preventing effective dissociation into individual cells and likely resulting in the mixed L/M cluster. S and UV cones, labelled for *arr3b*, and L/M cones labelled using *arr3b* via *in situ* HCR, showed similar expression patterns of the arr3 orthologs as in zebrafish (Renninger et al., 2011). Two major horizontal cell classes were delineated, namely *isl1-*positive *and lhx1a-*positive horizontal cells (Figure 4D, E). *in situ* HCR for *barhl2*, a gene specific to the *lhx1a*-positive horizontal cell subcluster, labelled horizontal cells as well as cells within the amacrine sublayer of the INL and the GCL (Figure 4F), as expected from our scRNA-seq dataset (Figure S6B). Subclustering of the bipolar cell population led to the identification of 26 clusters, highlighting the heterogeneity of this cell type within the killifish retina (Figure 4G, H). Labelling of cohorts of bipolar subclusters using *nxph1* and *dmbx1b* showed an apical to basal localisation within the INL, respectively (Figure 4I). Finally, subclustering of the amacrine cell population revealed four major amacrine types: glycinergic, GABAergic/parvalbumin, GABAergic, and starburst (Figure 4J, K). These amacrine cell types were spatially validated using *in situ* HCR for *slc6a9, pvalb5, gad2,* and *chata,* respectively (Figure 4L). Additional markers for glycinergic amacrines (*lamp5, tcf4*) are provided in Figure S5E. Interestingly, a visually appreciable difference in cell density could be observed for the amacrine subtypes, where parvalbumin and starburst amacrine cells seem less prevalent compared to GABAergic and glycinergic amacrines. Of note, we were also able to identify displaced starburst amacrines in the GCL (Figure 4L, arrowhead).

**Figure 4.**
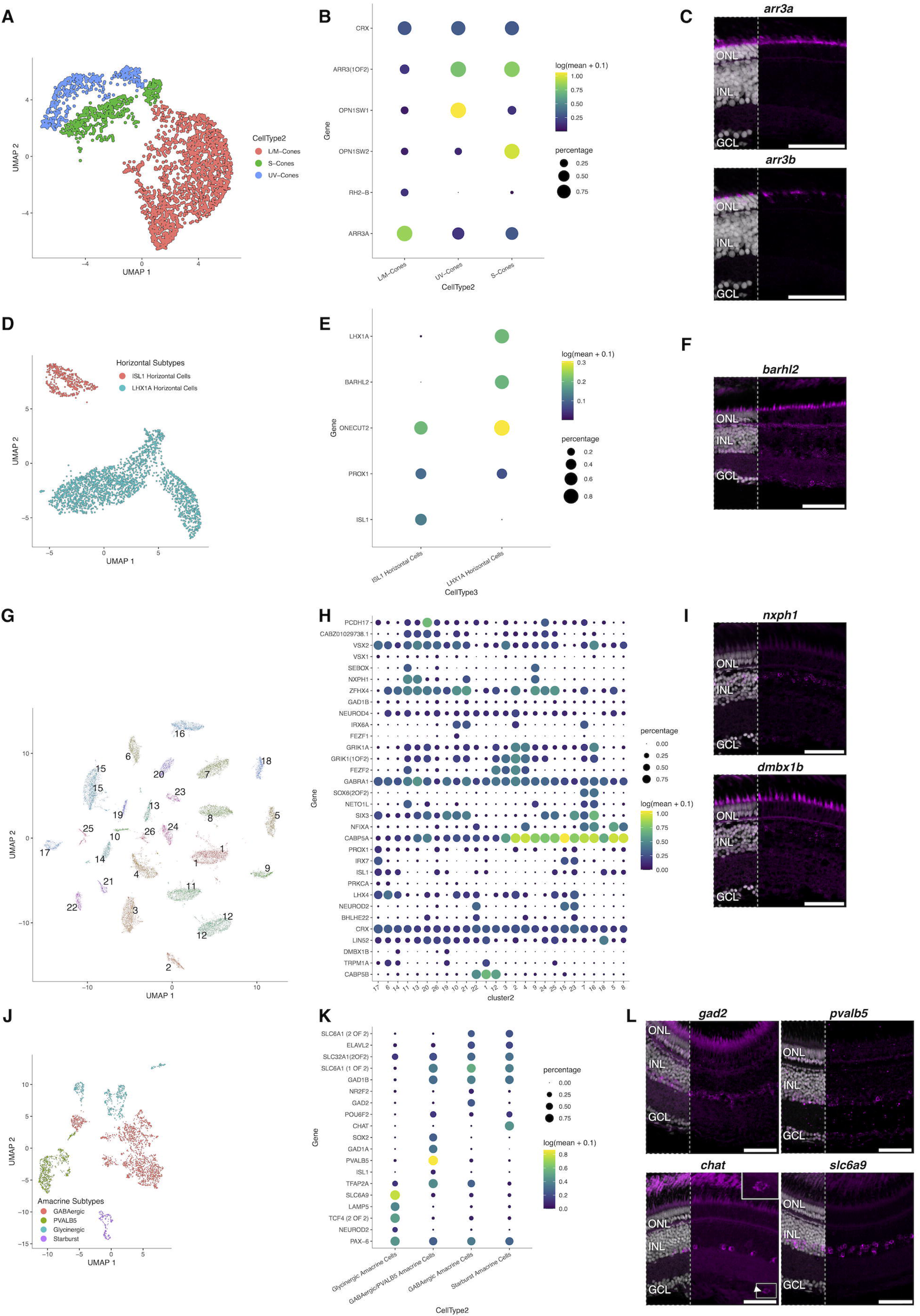
Validation of cell subtypes for specific killifish retinal populations. UMAP dimension reduction shows the subclustering of the photoreceptors (A), horizontal cells (D), bipolar cells (G) and amacrine cells (J). Cell type specific markers are shown as dot plots for each cell type (B, E, H and K, respectively). Dot size shows the percentage of cells expressing the marker gene while the colour indicates the mean transcript expression. *in situ* HCRs of subtype markers for photoreceptors (C), horizontal cells (F), bipolar cells (I), and amacrine cells (L) distinguish specific retinal cell subtypes. A displaced starburst amacrine (*chat+*) is highlighted with an inset box and arrowhead in the GCL (L). Images are acquired as mosaic Z-stack and are visualised as maximum projections. Merged images with nuclei shown in left third of image. Scale bars = 50 µm. GCL = ganglion cell layer, INL = inner nuclear layer, ONL = outer nuclear layer.

In conclusion, we have generated the first molecular survey of the cellular heterogeneity of the killifish retina, identifying all major cell classes typical of a vertebrate retina.

### 2.3| Integration of bulk and single-cell RNA-seq reveals cellular specificity and age-related dysregulation of retinal gene expression

We confirmed the robustness of the age-dependent transcriptomic signatures across RNA profiling techniques through comparisons of gene expression changes from the bulk RNA-seq within the scRNA-seq data (Figure S7). The up-regulated and unchanged gene sets detected by bulk RNA-seq showed similar age-dependent expression changes in the scRNA-seq dataset, although this was not observed for the down-regulated genes (Figure S7A, C).

Next, we sought to examine the cell-type specificity of the age-dependent differentially expressed transcripts identified in our bulk RNA-sequencing experiments through integration of the bulk and scRNA-seq dataset (Figure 5A, left; Figure S8A, left). A significant fraction (more than half) of the age-dependent, down-regulated transcripts from the bulk RNA-seq analysis displayed enriched expression within RPE cells (Figure S8). For the up-regulated genes, nearly half of the genes showed glial/immune enrichment (Figure 5A, left), this is notable as these cell types represent less than a third of the cells in the single-cell dataset and in the retina (Figure S2E). The aggregate of transcripts that increased expression with ageing were most highly expressed in glia and immune cells (Figure 5B).

**Figure 5.**
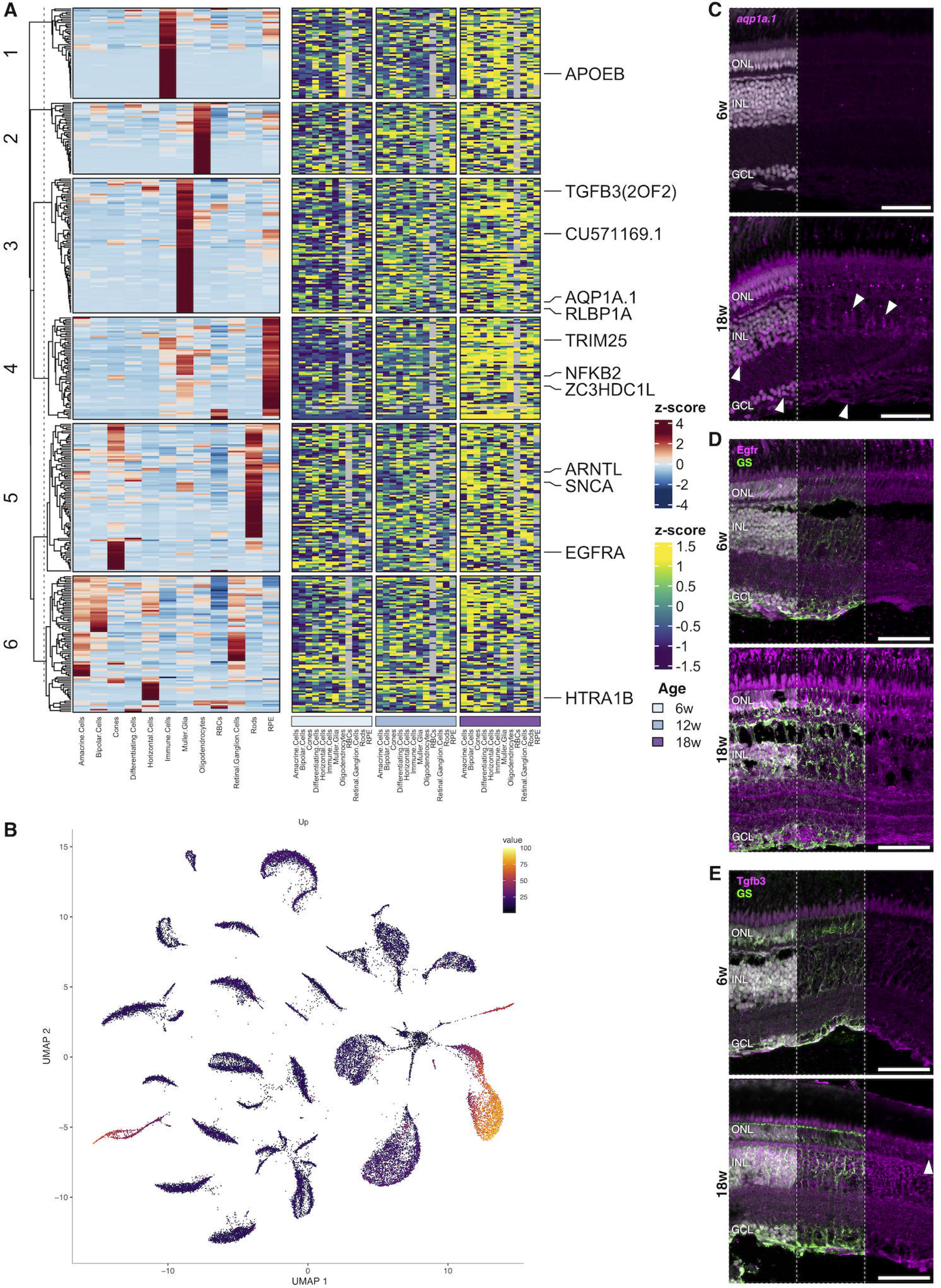
Killifish display age-associated transcriptional dysregulation, with many genes becoming expressed in all cell types. (A) Heatmaps of data integrated from both the bulk and scRNA-seq. Genes that are increasing with age in our dataset are typically expressed in a variety of cell types within the retina, with roughly half of the transcripts displaying enriched expression within glial/immune cells (left). With age, the genes are increasing in expression across numerous cell types (right). Several genes of interest are highlighted. ZC2HDC1L is *optn* and CU571169.1 is *slc7a11.* (B) UMAP showing that the genes that show increased expression with age are overall most highly expressed in Müller glia, immune cells, and oligodendrocyte clusters. (C-E) Spatial validation of up-regulated genes; total merge with nuclei shown in left third of image. (C) The Müller glia-specific gene *aqp1a.1* has transcript detected in many cell types at 18 weeks in in situ HCR, including retinal ganglion cells and photoreceptors. (D) Immunolabelling for Egfr and GS showing that Egfr protein is primarily in Müller glia at in young retinas – however, in old retinas there is increased expression in the ganglion cell and nerve fibre layers. Middle: Egfr + GS merge without nuclei; right: Egfr labelling only. (E) Antibody labelling for Tgfb3 and GS shows that Tgfb3 protein is restricted to Müller glia in young retinas but appears more universally expressed in old retinas, including the outer nuclear layer. Middle: Tgfb3 + GS merge without nuclei; right: Tgfb3 only. Images are acquired as mosaic Z-stack and are visualised as maximum projections. Scale bars = 50 µm. GCL = ganglion cell layer, GS = glutamine synthetase, HCR = hybridisation chain reaction, INL = inner nuclear layer, ONL = outer nuclear layer, RBCs = red blood cells, RPE = retinal pigment epithelium, w = weeks.

As our scRNA-seq dataset is age-matched to bulk RNA-seq samples, we plotted the relative expression of each gene within each cell type across time (Figure 5A, right; Figure S7A, right). Strikingly, for genes that showed expression highly enriched within specific cell types when evaluating the dataset as a whole (Figure 5A, left), we observed increased relative expression within many, if not all, retinal cell types of the aged killifish retina (Figure 5A, right). One prominent example was *trim25.* While its general expression is characteristic of immune and RPE cells (Figure 5A, left), in aged individuals all retinal cell types showed increased *trim25* transcript level as compared to their young counterparts (Figure 5A, right). Similarly, we observed that *snca* displayed enriched expression in rod photoreceptors (Figure 5A, left), but during ageing, relative *scna* expression increased in all other retinal cell types compared to their younger counterparts (Figure 5A, right). The Müller glia-specific gene *rlbp1a* (Figure 5A, left) displayed major dysregulation upon ageing, with an augmented expression in cones, differentiating cells, horizontal cells, immune cells, Müller glia, oligodendrocytes, and rods, but decreased relative levels in amacrine and bipolar cells across age (Figure 5A, right). Conversely, *arntl* expression did not show a clear confinement to a single retinal cell type (Figure 5A, left), but displayed increased expression in amacrine cells, bipolar cells, cones, immune cells, and oligodendrocytes upon ageing. Notably, this transcriptional dysregulation shown for many of the upregulated transcripts was not prominently observed for down-regulated genes (Figure S8A, right).

To validate the observed age-dependent spatial alterations of genes across cell types, we performed *in situ* HCR or immunohistochemistry to examine changes in localisation of selected genes. Aqp1 is associated with the oxidative stress response and the *aqp1a.1* transcript was generally expressed highest in Müller glia compared to the other cell types (Figure 5A, left). Upon ageing, transcript levels increased in Müller glia, but also in amacrine cells, differentiating cells, retinal ganglion cells, and oligodendrocytes (Figure 5A, right). In agreement, *in situ* HCR did not show appreciable *aqp1a.1* signal in 6-week-old retinas, presumably due to low expression, but in aged fish *aqp1a.1* expression appeared to be likely present in Müller glia as well as in amacrine or retinal ganglion cells in the INL and GCL (Figure 5C). Egfr is known to promote glial and progenitor cell proliferation (Yoo et al., 2021), but its transcript was overall most highly expressed in cone photoreceptors in our scRNA-seq dataset (Figure 5A, left). Upon ageing, *egfr* transcripts showed increased levels in cones, differentiating cells, horizontal cells, Müller glia, and oligodendrocytes (Figure 5A, right). Immunohistochemistry for Egfr revealed induced protein expression across all retinal layers in aged retinas (Figure 5D), confirming our scRNA-seq data. This dysregulated expression pattern was, however, not consistent for all genes. *tgfb3,* for instance, which was most enriched within Müller glia (Figure 5A, left), did not show a global increase in transcript expression across the retinal cell types, but rather a specific increase in rod photoreceptors and immune cells (Figure 5A, right). Nevertheless, Tgfb3 protein expression appeared to globally increase in aged killifish retina, including in the photoreceptor layer (Figure 5E). This discrepancy between our transcriptome data and immunohistochemical assessment might be due to transcriptional-translational decoupling, which has been reported in the killifish brain (Sacramento et al., 2020).

Altogether, our findings indicate that ageing killifish have a generalised loss of transcriptional homeostasis.

### 2.4| Single-cell RNA-seq shows cell type-specific transcriptome changes with age

To gain further insight into the age-associated changes in transcriptome expression at the level of individual cell types, we performed differential expression analysis across the ageing time points within the major retinal cell types. We utilized the regression modelling in Monocle3 to determine transcripts that changed with age (with cut-off: q-value < 1E-5). While single-cell differential analysis can have high false-positive rates (Squair et al., 2021), we noted interesting cell type-specific trends within the scRNA-seq dataset (Table S3). The changes observed below are between 6- and 18-week-old animals.

For example, we noted a decrease in transcript levels of the *atp1b3b* gene in retinal ganglion cells (log2FC = −1.008, q-value = 3,01E-13). This gene encodes the sodium/potassium-transporting ATPase subunit beta family b enzyme, which is essential in maintaining the electrochemical gradient across the cell membrane. Although ATP1B3 has not been directly linked to glaucoma, dysregulation of ion channel activity is known to disrupt ion balance within the retinal ganglion cells (Boal et al., 2023; Chen et al., 2013), leading to oxidative stress and apoptosis, both present in glaucomatous eyes (Ma et al., 2022). The decreased *atp1b3b* expression observed in aged killifish ganglion cells might thus indicate loss of cellular homeostasis and dysregulation of ion channel activity in retinal ganglion cells.

Interestingly, glial (Müller glia and oligodendrocytes) and immune cell populations exhibited similar transcriptional responses to age-related stress. Indeed, we observed increased transcript expression of *fkbp5* in Müller glia (log2FC = 1.257, q-value = 9.75E-80), immune cells (log2FC = 1.083, q-value = 1.44E-7), and oligodendrocytes (log2FC = 1.160, q-value = 2.99E-10). FKBP5 is an important modulator of stress responses through its binding to heat-shock protein 90 (HSP90), and increased levels of FKBP5 have been described to lead to a prolonged stress response (Zannas et al., 2015). Notably, Müller glia also showed increased levels of *hsp90aa1.2* (log2FC = 0.497, q-value = 1.07E-8), another gene linked to stress responses (Zannas et al., 2015).

Together, these data indicate that ageing-associated transcriptional dysregulation occurs across all cell types. We observe that numerous genes that function as stress response genes within glial and immune cell populations display upregulated expression, suggesting an influence for these cell types and transcripts for homeostatic management of stress responses or direct contributions to ageing phenotypes.

## 3| Discussion

The aim of this study was to identify the molecular changes in the ageing killifish retina that underlie retinal ageing/disease phenotypes by utilizing transcriptomic approaches. We observed age-related transcriptome changes between young (6-week-old), middle-aged (12-week-old), and old (18-week-old) female killifish retinas. Male killifish retinas were not included in this study, thereby excluding identification of potential sex-related transcriptional differences across ageing and the potential gender role during disease manifestation.

Our investigations revealed that the ageing killifish retina has dysregulation in pathways associated with oxidative stress, gliosis, and inflammageing, which were previously reported as hallmarks of brain ageing (Azam et al., 2021; Mattson and Arumugam, 2018). Additionally, increases in oxidative stress and inflammatory responses were corroborated by comparison to the ageing mouse retina, which had enrichment of similar cellular pathways (Xu et al., 2022). Interestingly, a previously published transcriptome study on the human and macaque retina also reported an age-related increase in oxidative stress and inflammation (Yi et al., 2021). In contrast to the aged murine retina (Xu et al., 2022) and a previous histological study of the aged killifish retina (Vanhunsel et al., 2021), we did not observe a substantial increase in senescence marker gene expression in killifish retinal cells at 18 weeks of age in our bulk RNA-seq dataset. This may be because previous killifish studies showed high levels of senescence-associated beta-galactosidase signal specifically in the RPE layer on histological preparations of 18-week-old retinas (Vanhunsel et al., 2021), a cell type that was removed during dissection in this study to focus on the neural retina. Further studies are required to understand the implications of senescence-related pathways and other potential important age-related changes of essential processes in RPE as it contributes to certain retinal disease, such as AMD (Somasundaran et al., 2020). Notably, minor increases in beta-galactosidase signal were previously detected in the neural retina of 24-week-old fish (Vanhunsel et al., 2021), suggesting the transcriptional dysregulation identified in our studies may precede senescence-associated phenotypes. Overall, determining the impact of genetic or pharmacologic manipulation of age-associated senescence, oxidative stress, and/or inflammation signatures on retinal health and function may open avenues for novel targets amenable for therapeutic intervention. Pharmacological manipulation of senescence using senolytic drugs has recently been shown to remove senescent cells from the aged killifish telencephalon, thereby restoring the neurogenic potential of the aged killifish brain (Van houcke et al., 2023). Similar pharmacological approaches targeting the identified ageing processes might thus have the potential to rejuvenate the aged retina.

We also observed transcriptional changes of several genes associated with age-related human retinal disease, which suggests that the killifish is a valuable translational model. Indeed, bulk RNA-seq revealed changes in specific genes, i.e., *optn, htra1b*, *and pcdh15a*, associated with glaucoma, AMD, and Usher syndrome, respectively (Ahmed et al., 2001; Rezaie et al., 2002; Yang et al., 2006). As increased levels of *optn* have been reported as neuroprotective (Markovinovic et al., 2018; Weil et al., 2018), the observed rise in *optn* expression in the aged killifish retina may suggest a compensatory phenotype attempting to attenuate age-related cellular stress. Altogether, our data highlight the value of old killifish retina as a model to study pathomechanisms of several human ageing retinal diseases. The killifish retina also has the potential for investigating age-related neurodegenerative brain pathologies (Bergmans et al., 2023a; de Bakker and Valenzano, 2023). For example, we observed that the expression of the Parkinson-associated gene, alpha-synuclein, is rising with age. Previous studies have reported a similar increase in the aged killifish brain that likely contributes to Parkinsonian-like phenotypes, such as protein aggregation and Lewy body formation (Bagnoli et al., 2022; Matsui et al., 2019). As the killifish is genetically amenable to transgenesis and CRISPR/Cas9 mutagenesis (Bedbrook et al., 2023; Hartmann and Englert, 2012; Rozenberg et al., 2023), it will be invaluable to perform genetic manipulation of disease genes within this rapidly ageing vertebrate.

Single-cell transcriptomics were used to delineate the cellular landscape of the killifish retina. The killifish retina contains all major cell classes, as well as oligodendrocytes, previously reported to be present in fish and avian retinas (Nakazawa et al., 1993; Santos-Ledo et al., 2023). Additionally, age-related changes in gene expression across the cell types were determined by combining the single-cell with the bulk RNA-seq datasets. In general, we observed age-related transcriptomic dysregulation within all retinal cell populations, showing that differentially expressed genes from bulk RNA-seq experiments displayed increased expression across all cell types upon ageing. Similar age-related transcriptional dysregulation has been described in *C. elegans* using a full organism single-cell atlas for a confined number of genes, although the mechanisms contributing to this altered gene expression remain elusive (Roux et al., 2023). One potential mechanism behind the changes in gene expression across various cell types includes a modified epigenome. Previous studies examining the retinas and RPE of patients presenting with AMD reported global changes in chromatin accessibility across both tissues (Wang et al., 2018). It will be interesting to determine the extent to which age-related transcriptomic changes in the killifish retina are mediated by alterations to the epigenetic landscape. Furthermore, it is important to note that RNA levels are not indicative of protein levels *in vivo*. Decoupling of transcription and protein synthesis has been observed in killifish and was shown to worsen with age (Sacramento et al., 2020). As such, the transcriptional changes reported here may not fully reflect the changes at protein level and may contribute to an underestimation in the severity of age-related retinal dysfunctional.

In summary, we have identified age-related gene expression dysregulation in the killifish retina which may be amenable for therapeutic intervention to prevent deleterious effects of ageing on visual function. This work also highlights that the rapidly ageing killifish can be further utilised to define the molecular features underlying age-related retinal pathology and vision loss. As the retina is considered a window to the brain, our data may provide insight into processes occurring within the ageing brain and demonstrate that the short-lived killifish is a unique animal model to investigate the consequences of molecular dysregulation on CNS health. This allows for the identification of means to prevent age-related loss of neural integrity and function.

## 4| Materials & Methods

### 4.1| Fish husbandry

Adult female African turquoise killifish of the *Nothobranchius furzeri* GRZ-AD inbred strain were used. Fish were bred in-house as described (Bergmans et al., 2023b; Vanhunsel et al., 2022, 2021). Briefly, killifish hatchlings were fed twice a day with brine shrimp (*Artemia salina* nauplii, Ocean Nutrition) during the first two weeks post hatching. Next, until adulthood (5-weeks-old), fish were fed with both brine shrimp and *Chrinomidae* larvae twice a day, after which adult fish were fed twice a day with only *Chironomidae* larvae. Hatchlings were reared in small (1 L) tanks for the first week (50% daily water renewal), after which they were housed in 3.5L tanks in a multilink ZebTec (Techniplast) system ensuring standardized conditions (temperature 28.3°C; conductivity 600 µS, pH 7; 12h light/dark cycle; water renewal 15%). Adult fish were always grouped with three female fish and one male fish. Males were removed from the age of 12-weeks-old due to overdominance. Based on previous in-house lifespan and biased ageing studies, three different age groups were selected, i.e., young adult (6-weeks-old), middle-aged (12-weeks-old) and old (18weeks-old) fish (Van houcke et al., 2021; Vanhunsel et al., 2021). The Ethical Committee for animal experimentation of KU Leuven, strictly following the European Communities Council Directive of 2010 (2010/63/EU) and Belgian legislation (Royal degree of 29 May 2013), approved all animal experiments.

### 4.2| Tissue collection/processing

#### 4.2.1| Tissue/sample collection and dissociation for (sc)RNA-sequencing

Fish were euthanized using 0.1% tris buffered tricaine (Merck) in system water and their body size was measured using callipers. Retinas were collected as previously described for zebrafish (Van Dyck et al., 2023). Briefly, the visible part of the dermal layer of the cornea, the scleral layer of the cornea and lens were removed, thereby exposing the retina. Using sterile Dulbecco PBS (DPBS, ThermoFisher Scientific), the retinal pigment epithelial was rinsed off as much as possible. Next, the retinas were collected (cut at the optic nerve head) and either snap frozen in liquid nitrogen for bulk RNA sequencing (RNA-seq) or collected in medium for the generation of single-cell suspensions to be used for single-cell RNA sequencing (scRNA-seq).

For the bulk RNA-seq experiments, both retinas of one fish were pooled as a single sample and 10 samples (fish) were used per age group. Retinal samples were digested using Tri-reagent (Sigma-Aldrich) before purification of the total RNA using the RNeasy kit (Qiagen) according to the manufacturer’s protocol and stored at −80°C until library preparation (Bergmans et al., 2023d). Just prior to library preparation, final RNA quality and integrity were assayed using the DNA 12000 kit on the Bioanalyzer (Agilent) at the KU Leuven Genomics Core (https://www.genomicscore.be/).

For the scRNA-seq experiment, single-cell suspensions were generated as described (Bergmans et al., 2023b). Briefly, both retinas from either three (6-week-old) or two (12- or 18-week-old) fish were pooled and collected in Leibovitz 15 (L15, ThermoFisher Scientific) medium complemented with 2% Penicilin-Streptomycin (ThermoFisher Scientific), 0.5% Gentamycin (Merck), 2% heat-inactivated Fetal Bovin Serum (ThermoFisher Scientific), 1% Glutamax (ThermoFisher Scientific), 2% MEM Essential Amino Acids (ThermoFisher Scientific), 2% B27 (ThermoFisher Scientific), 25 mM D-Glucose (Merck), 25 mM HEPES (ThermoFisher Scientific) and brought to pH7.4, referred to as complete L15. Retinas were rinsed three times with complete L15 before enzymatic digestion using sterile activated papain (16 U/mL, Worthington) for 30 min at 28°C. Next, digested retinas were rinsed three times with complete L15. Single-cell suspensions were obtained after mechanical trituration (10 times with both a p1000 and a p200 pipet) in sterile PBS with 1% bovine serum albumin (fraction V, Sigma-Aldrich). To ensure singlets, cell clumps were removed using a 40 µm cell strainer (PluriSelect). Cell viability was measured with the LUNA Automated Cell Counter at the KU Leuven Genomics Core. Two biological samples, from independent animals were processed at each age, totaling six scRNA-seq runs.

#### 4.2.2| Tissue collection for in situ hybridization, immunohistochemistry and imaging

Fish were euthanized as described above. Intracardial perfusion with phosphate buffered saline (NaHPO_4_.2H_2_O, 8mM; KH_2_PO_4_, 2 mM; NaCl, 0.15M; KCl, 3mM; pH 7.4) and 4% paraformaldehyde (PFA, in PBS, Merck-Aldrich) was executed, all as described (Bergmans et al., 2023c; Mariën et al., 2022). Eyes were collected and post-fixed overnight in 4% PFA, followed by three rinses in PBS. Next, eyes were incubated until saturation using an increasing series of sucrose concentrations (10, 20, 30% weight by volume in PBS) and embedded in 1.25% agarose and 30% sucrose in PBS for cryosectioning. For each eye, 10 µm sagittal cryosections were serially collected on eight SuperFrost Plus Adhesion Slides (Epredia) and stored at −20°C until further use (Bergmans et al., 2023c).

### 4.3| RNA-sequencing

#### 4.3.1| Bulk RNA-seq data analysis

Libraries were generated using the QuantSeq 3’ (Lexogen) mRNA protocol and subsequently sequenced using the Illumina HiSeq 4000 (Illumina; 50bp single end reads). On average 8.2M reads were obtained for each of the 10 replicate samples collected from each age (6-, 12- and 18-week-old fish). Reads were trimmed using Trim-Galore! (v0.6.7), a wrapper for Cutadapt (v3.4) and FastQC (v0.11.9). Trimmed reads were mapped to a custom-built transcriptome (see below) using STAR (v2.7.0) (Dobin et al., 2013). Mapped reads were cleaned using Samtools (v1.9.4), gene counts generated by HTseq (v0.11.2) (Anders et al., 2015), and visualizations generated in R (v3.6.1). Comparative analysis between individual ages was performed using EdgeR (v3.42.4) (Robinson et al., 2010). Thresholds for statistical significance were set at |log2FC| ≥ 1 & FDR ≤ 0.05. As the Killifish reference genome is still in a draft phase, gene annotation for those passing statistical thresholds were checked by hand and in some instances re-annotated to conform to the nearest ortholog using BLAST (Altschul et al., 1990). Heatmaps were generated using ComplexHeatmap (Gu et al., 2016) and PCA using the R stats package while GO was performed using the Gene Ontology knowledgebase resources (Ashburner et al., 2000; Consortium et al., 2023).

#### 4.3.2| 10X Genomics sequencing, and data analysis

Single-cell suspensions with a viability score greater than 85% (ranging from 87.20% to 95.00%) were loaded on to the 10X Genomics microfluidic system with a capture target of 10,000 cells per sample. scRNA-seq was performed using standard protocols for 10x Genomics v3 3’ sequencing. Samples were sequenced to a target read depth of ∼160 million paired end reads per library (19,291 reads/cell on average) using Illumina NovaSeq (Illumina). Library preparation and sequencing were performed at the KU Leuven Genomics Core (https://www.genomicscore.be/).

To increase mapping efficiency of sequencing read, we built a custom transcriptome, utilizing the reference sequence (GRZ Assembly, 05/2015) (Valenzano et al., 2015) and custom annotations. Using GffCompare tool, the annotations were customized from Cellranger ‘mkgtf’ by combining the NCBI annotation file (NFINdb) with the in-house sequenced PACBIO-ISOSEQ annotation files for killifish telencephalon. This way the gtf files from PACBIO long-reads and the NFIN db reference transcripts were efficiently merged (Rajagopal et al., 2021). This helped us improve our final genomic annotation file and ensured higher mapping accuracy and coverage for performing single-cell RNA-seq analysis. Use of the custom transcriptome increased mapped reads by 11%, resulting in a 9% increase in cell number compared to the standard reference transcriptome alone. Sequencing reads were mapped to the custom transcriptome using CellRanger-7.0.1.

Mapped scRNA-seq data was processed in Monocle3 (monocle3_1.2.9) (Cao et al., 2019; Trapnell et al., 2014). Cells utilized for analysis contained greater than 1,000 unique transcripts and less than 20,000 transcripts. UMAP dimension reduction was performed on 3,327 high variance genes, determined from the transcripts/10,000 transcripts normalized count matrix, with the top 22 principal components used as input into the Monocle3 reduce_dimension function (max_components = 2, reduction_method = “UMAP”, preprocess_method = “PCA”, umap.metric = “euclidean”, umap.min_dist = .25, umap.n_neighbors = 25, build_nn_index = TRUE). Cell type assignment of clusters was performed using known markers of vertebrate retinal cell types, all as performed previously (Clark et al., 2019; Lu et al., 2020). Further subclustering of individual cell types (Amacrine Cells, Horizontal Cells, Cones, Bipolar Cells) was performed to examine marker genes of putative cellular subtypes.

### 4.4| Immunohistochemistry, in situ hybridisation chain reaction, and imaging

Immunohistochemistry (IHC) or *in situ* hybridization chain reaction (HCR) were utilised to identify spatial expression patterns of targets of interest from transcriptional profiling. In preparation for IHC, slides were warmed to room temperature and tissue rehydrated with PBS. Antigen retrieval with sodium citrate pH 6.0 was performed by boiling in coplin jars for 20 minutes. Cooled slides were then washed with PBS and placed in blocking solution (10% normal goat serum, 1% BSA in PBS) for 1 hour. Primary antibodies were mixed in blocking solution, then applied to the slides overnight at 4°C. Primary antibody was washed off with PBS thrice, then secondary antibodies mixed 1:1000 in blocking solution were incubated overnight at 4°C. After rinsing with PBS three times, slides were mounted and imaged. A list of antibodies and their respective concentrations is available in Table S4.

Fluorescent in situ mRNA visualization was accomplished using *in situ* HCR v3.0, following previously described protocols (Choi et al., 2018; Elagoz et al., 2022; Van houcke et al., 2021). Table S5 provides a list of target genes and probe pair design. Briefly, HCR v3.0 probes sequence targets were obtained using Easy_HCR (https://gitlab.com/NCDRlab/easy_hcr) and probe pools were ordered from Integrated DNA Technologies (IDT). Cryosections were dried at 37°C, rehydrated using diethyl pyrocarbonate (DEPC, Acros) -treated PBS (0.1% v/v), and sections were permeabilized using proteinase K (10 µg/mL in PBS-DEPC, Roche) for 10 min at 37°C. Permeabilization was stopped by incubation in 4% PFA for 10min at room temperature. Slides were rinsed three times before incubation in hybridization buffer (formamide, 30%; NaCl, 0.75M; sodium citrate, 75mM; citric acid pH6.0, 9 mM; Tween20, 0.1%; Heparin, 50 µg/mL; Denhardt’s solution 50x, 2%; Dextran, 10%) for 30 min at 37°C. Probes were added at a final concentration of 6.67 nM in probe hybridization buffer and hybridization was accomplished overnight at 37°C. Slides were rinsed three times and incubated with amplification buffer (NaCl, 0.75M; sodium citrate, 75 mM; Tween20, 0.1%; Dextran, 10%) for 30 min at room temperature. Both hairpins to visualize a probe set were prepared separately: 3pmol of each hairpin (H1 & H2) was incubated for 90s at 95°C and cooled down to room temperature for 30 min. Hairpins were added to the slides at a final concentration of 40 nM in amplification buffer and incubated overnight at room temperature. Nuclear labelling was achieved using 4’,6-diamidino-2-phenylindole (DAPI), prior to tissue mounting using Mowiol®.

All retinal sections were imaged on a Zeiss LSM900 microscope with a Plan-Apochromat 20X 0.8 NA objective in Airyscan CO-2Y mode. Images were acquired as mosaic Z-stacks covering the whole thickness of the section and visualized as maximum projections.

## Data Availability

GEO accession number is under review and will be made publicly available upon final publication.

## Acknowledgments

We thank Simon Buys and Arnold Van den Eynde for the daily fish care and environmental control. Animal housing/maintenance was supported by a KU Leuven small equipment grant KA-16-00745. The research was financially supported by Research Foundation Flanders (FWO Vlaanderen, Belgium; G092222N) and KU Leuven Research Council (C3/21/012#56343072). SB and LM hold a Research Foundation Flanders (FWO Vlaanderen, Belgium) PhD fellowship (1S6617N and 1S42720N, respectively). NCLN was supported by a BrightFocus Macular Degeneration Fellowship. AMK was supported by a Moorfields Eye Charity PhD studentship (GR001503) to RBM. RBM was supported by a BBSRC David Phillips Fellowship (BB/S010386/1), and a BBSRC Partnering award (BB/V018078/1) to RBM, PAR, BSC and C-KH. C-KH was supported by National Institutes of Health DP2AG077431. PAR and BSC were supported by an unrestricted grant to the Department of Ophthalmology and Visual Sciences at Washington University School of Medicine and individual career development awards from Research to Prevent Blindness and by the National Eye Institute of the National Institutes of Health under award number P30EY002687.

## Author contributions

Conceptualisation & experimental design: SB, NCLN, LuM, LA, PAR, RBM, BSC, LM. Data curation: SB, NCLN, AMK, JDDS, RBM. Bioinformatic analysis: EH, C-KH, PAR, BSC. Reference genome: RA, LA. Visualisation: SB, NCLN, PAR, BSC. Writing – original draft preparation: SB, NCLN. Reviewing & editing: SB, NCLN, LuM, EH, AMK, JDDS, RA, C-KH, LA, PAR, RBM, BSC, LM. Funding acquisition: SB, NCLN, LuM, RBM, LM. Project administration: SB, NCLN, PAR, BSC, RBM, LM. Supervision: PAR, BSC, RBM, LM.

## Conflict of interest

The authors declare that they have read and approved the manuscript and have no conflicts of interest.

---

**Figure S1. Heatmap showing differential transcript expression across all bulk RNA-seq samples.** Differentially expressed genes were identified using the following thresholding criteria: FDR < 0.05 and |log2FC| ≥ 1. Over age, biological replicates are highly consistent, observed changes are gradual with time, and killifish show more up-regulated genes compared to down-regulated genes. w = weeks.

**Figure S2. Quality control of single-cell RNA-sequencing.** UMAP dimension reductions indicating the (A) log10(Total_mRNAs) or (B) cluster designation for each cell. (C) Graph depicting the sample proportions for cells within each cluster. (D) UMAP dimension reductions highlighting the presence of cells from each age (6, 12, and 18 weeks) within all clusters. (E) Cell type proportions of retinal cells within each sample. (F) Boxplot showing the average transcript count across cells within each sample. Overlayed dots show individual transcript counts for each cell, with cells coloured by the annotated cell type. (G) Boxplots highlighting the transcript counts of cell types across the profiled ages, highlighting consistency of transcript counts within cell types across the ages, but variability across individual cell types. RBC = red blood cell; RPE = retinal pigment epithelium.

**Figure S3. Cell type marker gene expression.** UMAP dimension reductions highlighting marker gene expression used for cell type calling across the dataset.

**Figure S4. Cell type markers for RBC and RPE cells.** (A) Dot plot of marker genes used to identify the RPE and RBC populations. The dot size shows the percentage of cells expressing a specific gene while the colour indicates expression level. (B) *in situ* HCR for *rpe65a* shows that RPE65a is confined to the RPE. Images are acquired as mosaic Z-stack and are visualised as maximum projections. Scale bar = 50 µm. GCL = ganglion cell layer, HCR = hybridisation chain reaction, INL = inner nuclear layer, ONL = outer nuclear layer, RBC = red blood cell; RPE = retinal pigment epithelium.

**Figure S5. Additional spatial validation for cell type marker genes via *in situ* hybridisation.** Additional cell type-specific markers from scRNA-seq for each retinal cell type are validated on cryosections using *in situ* HCR. All photoreceptors are labelled by *crx* (A) while rods are specifically labelled by *nrl* (B). Horizontal (C) and bipolars (D) cells are shown by *in situ* labelling for *tfap2a* and *otx2*, respectively. (E) All amacrines are labelled by the canonical amacrine marker *slc32a1b,* while glycinergic amacrine cells are labelled with *lamp5* and *tcf4*. (F) Retinal ganglion cells are shown by *slc17a6b* labelling. (G) Differentiating cells are shown in the ciliary marginal zone at the periphery of the retina by expression of *stmn1a* and *fgfbp3.* First panel for both markers is merged with nuclei stain. Second panel is marker only. (H) Characteristic radial fibre labelling for Müller glia is accomplished by HCR for *rlbp1a,* while low expression of *gfap* is detected in Müller glia endfeet. (I) *in situ* HCR for *cldn19* shows the sparse presence of oligodendrocytes in the GCL and INL (arrowhead), while a high density is observed in the optic nerve head. Images are acquired as mosaic Z-stack and are visualised as maximum projections. (A-F, H) Merged images with nuclei shown in left third of image. Scale bars = 50 µm. GCL = ganglion cell layer, HCR = hybridisation chain reaction, INL = inner nuclear layer, ONL = outer nuclear layer.

**Figure S6. Heatmaps showing scRNA-seq expression of cell subtype marker genes across all cell types identified.** (A) Cone subtype marker genes; (B) Horizontal cell subtypes; (C) Bipolar cell subtypes; (D) Amacrine cell subtypes. Coinciding with the *in situ* expression pattern observed in Figure 4, *barhl2* (B) transcript is detected most abundantly in horizontal cells, but is also observed in ganglion and amacrine cells.

**Figure S7. Comparison of age-related transcriptional changes between the bulk and single-cell RNA-sequencing datasets.** Boxplots indicating the age-dependent fold change of differentially expressed genes from bulk RNA-seq within the scRNA-seq dataset, comparing (A) Fold-change of expression comparing 12-week and 6-week-old retinas across all cell types, or (B) cell type-specific changes across those ages. (C) Boxplots indicating transcript fold change across all cell types, or (D) within individual cell types in 18-week-old retinas in comparison to 6-weeks-old fish. On average, genes displaying differential expression by bulk RNA-seq analysis displayed corresponding changes within the scRNA-seq dataset.

**Figure S8. Heatmap and UMAP for genes with reduced expression between young and old killifish retinas.** (A) Heatmap of integrated bulk and scRNA-seq, showing the cell types that normally express those genes (left) and the changes in expression across the ages investigated for the cell types (right). (B) UMAP showing that the genes that are going down in expression with age are typically expressed in the RPE. There are several bright cells within the RPE cluster; see inset. RBCs = red blood cells, RPE = retinal pigment epithelium.

**Table S1. List of all genes picked up during bulk RNA-sequencing across age (6-, 12- and 18-week-old) including XLOC ID#s.** Differentially expressed genes were identified using following cut-offs: FDR < 0.05 and |log2FC| ≥ 1. Up-regulated genes are highlighted in orange, down-regulated genes are in blue, and unchanged are not highlighted. The table shows the gene XLOC ID#, two killifish gene short names, orthologue gene in zebrafish and human after nucleotide BLAST as well as its query cover and E-value. Raw data are depicted for every biological replicate as well as the cpm. Furthermore, log2(FC), p-values and FDRs are provided for all inter-age comparisons. cpm = count per million, FC = fold change, FDRs = false discovery rates.

**Table S2. Detailed list containing all GO-categories of up- and down-regulated biological processes and molecular functions.** Sheet one shows all the up-regulated biological processes (BP) after GO-term analysis. Sheet two shows all the down-regulated BP. Up-regulated molecular functions are depicted in sheet three while down-regulated MF are shown in sheet 4. FDR = false discovery rate.

**Table S3. List of all differentially expressed genes per cell type across age (6-, 12-, and 18-week-old) identified by scRNA-seq.** Regression modelling in Monocle3 determines transcripts that change with age (with cut-off criteria: q-value < 1E-5). Mean expression for each age is listed, including the Age (term column) at which statistical significance was determined. Differential expression within the full data or differential expression within individual cell types are listed as separate tabs of the spreadsheet. Exprs = expression, FC = fold change.

**Table S4. List of antibodies used for immunohistochemistry.**

**Table S5. List of probe pairs used for in situ hybridisation chain reaction experiments.** A list containing all the genes that were spatially validated using *in situ* hybridisation chain reaction. The list contains the NCBI gene IDs as well as the corresponding RefSeq association codes. The number of probe pairs per gene are shown as well as the amplifier(s) that were used.

## Notes

### Competing Interest Statement

The authors have declared no competing interest.

### Summary of Updates

Figure 1 revised (gene # on diagram); zoom boxes added on Figure 2D; Figure 3C plot updated to include GNAT1; zoom box added to Figure 4L.

